# De novo serine biosynthesis from glucose predicts sex-specific response to antifolates in non-small cell lung cancer cell lines

**DOI:** 10.1101/2021.12.22.473923

**Authors:** Jasmin Sponagel, Siddhartha Devarakonda, Joshua B. Rubin, Jingqin Luo, Joseph E. Ippolito

## Abstract

Lung cancer is the leading cause of cancer-related death. Intriguingly, males with non-small cell lung cancer (NSCLC), the most common type of lung cancer, have a higher mortality rate than females. Here, we investigated the role of serine metabolism as a predictive marker for sensitivity to the antifolate pemetrexed in male and female NSCLC cell lines. Using [^13^C_6_] glucose tracing in NSCLC cell lines, we found that male cells generated significantly more serine from glucose than female cells. Higher serine biosynthesis was further correlated with increased sensitivity to pemetrexed in male cells only. Concordant sex differences in metabolic gene expression were evident in NSCLC and pan-cancer transcriptome datasets, suggesting a potential mechanism with wide-reaching applicability. These data were further validated by integrating antifolate drug cytotoxicity and metabolic pathway transcriptome data from pan-cancer cell lines. Together, these findings highlight the importance of considering sex differences in cancer metabolism to improve treatment for all patients.

## Introduction

Lung cancer is the second most common cancer and the leading cause of cancer related death in the U.S. and worldwide (Siegel et al., 2021; Sung et al., 2021). Non–small cell lung cancer (NSCLC) is the most common subtype, accounting for approximately 80%–85% of all lung cancers (Howlader et al. 2021). Most patients with NSCLC are diagnosed at advanced stages and generally have a poor prognosis with a median survival of 8-10 months (Shih et al., 2020; Siegel et al., 2021). One of the front-line chemotherapeutics for patients with non-squamous NSCLC is pemetrexed (Ettinger et al., 2021), an antifolate chemotherapeutic. Although the addition of pemetrexed, either as a first-line treatment in combination with carboplatin or cisplatin or as a monotherapy for second-line treatment, has improved survival and overall response rate in men and women, about 50% of advanced NSCLC patients show no response to treatment (Ettinger, 2002; Jo et al., 2018; Park et al., 2015; Shih et al., 2020). Thus, better biomarkers for treatment response are needed to improve survival for all patients with NSCLC.

As in most cancers, males with NSCLC exhibit a higher mortality rate compared to females (Dong et al., 2020; Moore et al., 2004; de Perrot et al., 2000; Siegel et al., 2021; Wainer et al., 2018). This sex difference in mortality may be, at least partially, driven by sex differences in response to the current treatment. Indeed, multiple phase 3 clinical trials have shown that female NSCLC patients respond better to certain treatment regimens, including pemetrexed-based treatment combinations (Paz-Ares et al., 2012; Wang et al., 2017). Pemetrexed inhibits multiple folate cycle enzymes, thus, inhibiting DNA synthesis and cell replication (Adjei, 2004). The folate cycle is a metabolic nexus of multiple metabolic and cellular processes, such as providing purines and pyrimidines for DNA synthesis, serine and glycine metabolism, thymine and methionine synthesis, and redox homeostasis (Yang et al., 2021). Whether the activity of these metabolic pathways could be utilized to predict sensitivity to antifolates, has yet to be determined.

Most metabolic pathways exhibit sex differences across many species from fertilization to adulthood (Bermejo-Álvarez et al., 2008; Van Blerkom, 2008; Bredbacka and Bredbacka, 1996; Gutiérrez-Adán et al., 2001, 2006; Hedrington and Davis, 2015; Krumsiek et al., 2015; Pergament et al., 1994; Ray et al., 1995; Rubin et al., 2020; Tagirov and Rutkowska, 2014; Tiffin et al., 1991). Thus, we hypothesized that sensitivity to pemetrexed may be driven by different metabolic pathways in male and female NSCLC. Previously, we provided support for this idea when we identified glycolytic transcriptome and metabolome signatures that selectively stratified male, but not female patients with lower grade gliomas (Ippolito et al., 2017). These data suggest that sex-specific analyses of predictive metabolic biomarkers may help stratify patients for treatment approaches.

Here, we provide further support for the presence of sex differences in cancer cell metabolism and its impact on predicting treatment response in male and female cancer cell lines. Using [^13^C_6_] glucose tracing datasets of over 80 NSCLC cell lines (Chen et al., 2019), we demonstrate that male cell lines have increased *de novo* serine and glycine biosynthesis from glucose compared to female cell lines. Higher serine biosynthesis correlated with increased sensitivity to pemetrexed in male NSCLC cell lines only. Concordant sex differences in metabolic gene expression were evident in NSCLC and pan-cancer transcriptome datasets, showing increased transcript enrichment of this pathway in male tumors. These findings were also validated with drug cytotoxicity and gene expression analyses of pan-cancer cell lines obtained from The Cancer Cell Line Encyclopedia (TCLE). In summary, we discovered sex differences in the role of *de novo* serine biosynthesis in male and female NSCLC cell lines, which may be leveraged to better predict sex-specific treatment response to antifolates in NSCLC patients. Thus, our data highlight the importance of considering sex, even in established cancer cell lines, as understanding sex differences in cancer metabolism and its impact on therapeutic response will lead to the development of new diagnostic and therapeutic approaches to improve cancer outcomes for both men and women.

## Results

### Male NSCLC cell lines utilize more glucose for *de novo* serine biosynthesis

Serine is a non-essential amino acid that can fuel the folate cycle by donating a one-carbon unit to the synthesis of glycine. It can be either taken up by the cell or synthesized *de novo* from glycolytic intermediates (Locasale, 2013). Upon its uptake into the cell, most glucose is converted to pyruvate via the glycolytic pathway, which can then be converted into acetyl-CoA to replenish the TCA cycle. However, glycolysis also provides 3P-glycerate, the substrate for serine biosynthesis, via the phosphoserine pathway (**Fig. 1A**). To determine whether male and female NSCLCs may differ in their requirement for *de novo* serine synthesis, we first investigated whether male and female NSCLC cell lines utilize glucose differently. We assessed published stable isotope nutrient tracing data of 57 male and 30 female NSCLC cell lines cultured in [^13^C_6_] glucose for either 6 or 24 hours (Chen et al., 2019). While total label enrichment was similar in male and female cell lines in TCA cycle metabolites we found that male cell lines incorporated more glucose into serine and glycine biosynthesis than female cell lines (**Fig. 1B-C**). Interestingly, density plots revealed an almost bimodal distribution in the male cell lines (particularly for serine), suggesting that the male population consists of a high and a low *de novo* serine and glycine biosynthesis population. The bimodal distribution in female cell lines was almost absent, indicating that the female population is primarily comprised of a low *de novo* serine and glycine biosynthesis population. Together, these data suggest that male NSCLC cells may have a greater requirement for glucose-dependent *de novo* serine and glycine biosynthesis.

**Figure 1:**
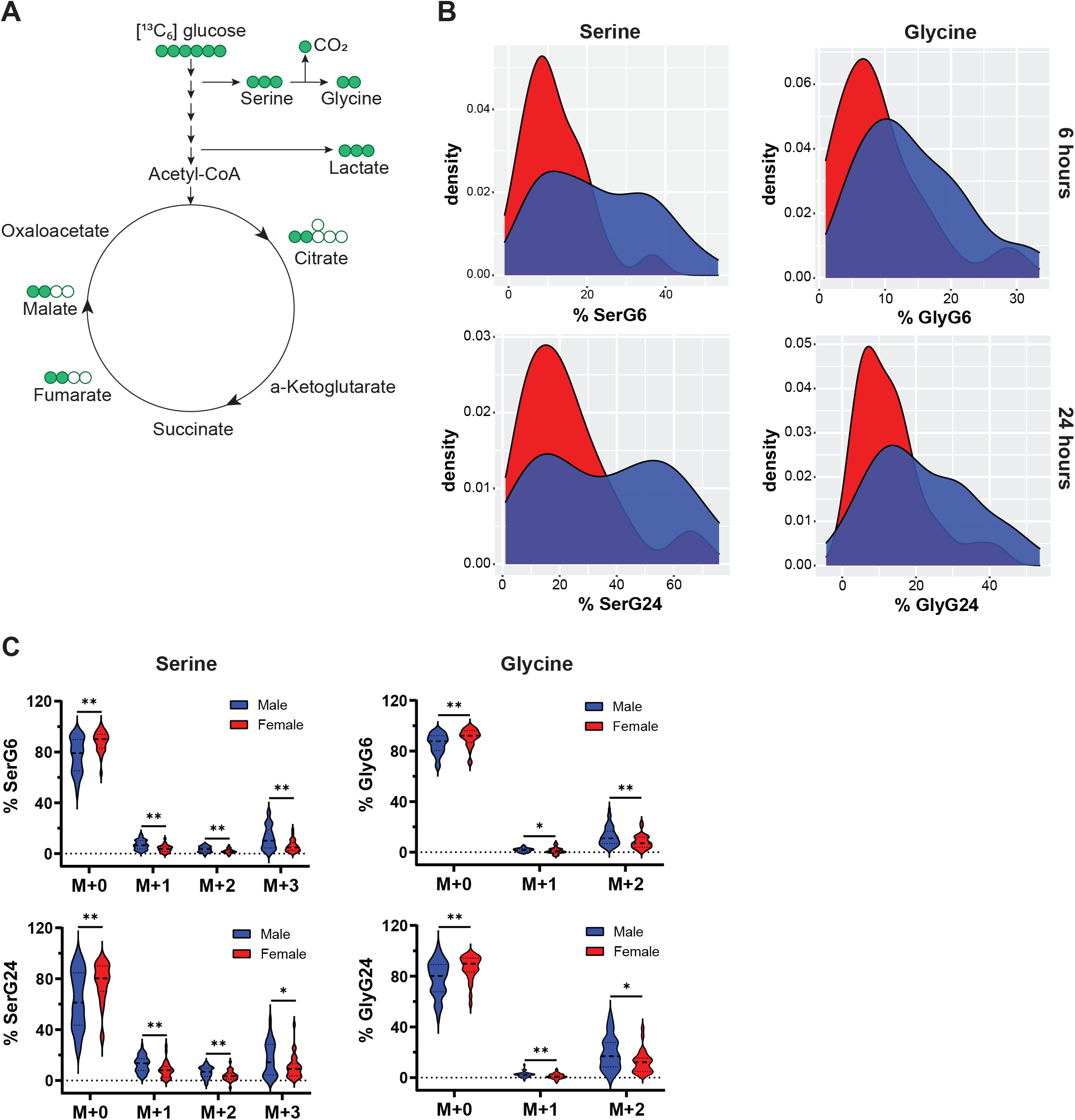
Male NSCLC cell lines utilize more glucose for *de novo* serine biosynthesis. A, Schematic of representative isotopologues generated from [^13^C_6_] glucose-based labeling. B, Density plots showing [^13^C_6_] glucose enrichment in serine and glycine after 6 and 24 hrs in male (n=49) and female (n=28) non-small cell lung cancer (NSCLC) cell lines. C, Violin plots showing isotopologue enrichment of [^13^C_6_] glucose into serine and glycine after 6 and 24 hrs in male (n=49) and female (n=28) NSCLC cell lines. All possible isotopologues are shown. Serine: M+0, M+1, M+2, M+3; glycine: M+0, M+1, M+2. *q<0.05, **q<0.01; Mann-Whitney test, q = FDR-adjusted p-values. Abbreviations: ser, serine; gly, glycine; G, glucose

### *De novo* serine biosynthesis from glucose is associated with sensitivity to antifolate drugs in male NSCLC cell lines

Serine is an important one-carbon donor to the folate cycle and therefore contributes to nucleotide synthesis, NADPH generation, and methylation reactions (Locasale, 2013; Minton et al., 2018; Mullarky et al., 2016; Singh et al., 2021; Tanaka et al., 2021; Tedeschi et al., 2013; Ye et al., 2014). The folate cycle is a well-studied target of multiple antifolate therapeutics. In fact, the antifolate pemetrexed is a front-line chemotherapeutic for patients with non-squamous NSCLC (Ettinger et al., 2021).

As the serine m+3 isotopologue had the highest correlation with pemetrexed when all of these cell lines were evaluated as a whole previously (Chen et al., 2019), we correlated label enrichment of serine m+3 and glycine m+2 with pemetrexed IC_50_ values in male and female cell lines separately. We found that sensitivity to pemetrexed and the combination of pemetrexed and the alkylating agent cisplatin, another front-line chemotherapeutic given with pemetrexed for NSCLC (Ettinger et al., 2021), correlated negatively with label enrichment in serine and glycine in male, but not female, NSCLC cell lines (**Fig. 2A-B**). Interestingly, no other chemotherapeutic agents tested showed a significant correlation in either the male or female cell lines. These data indicate that increased *de novo* serine and glycine biosynthesis in male NSCLC cell lines predicts increased sensitivity to pemetrexed.

**Figure 2:**
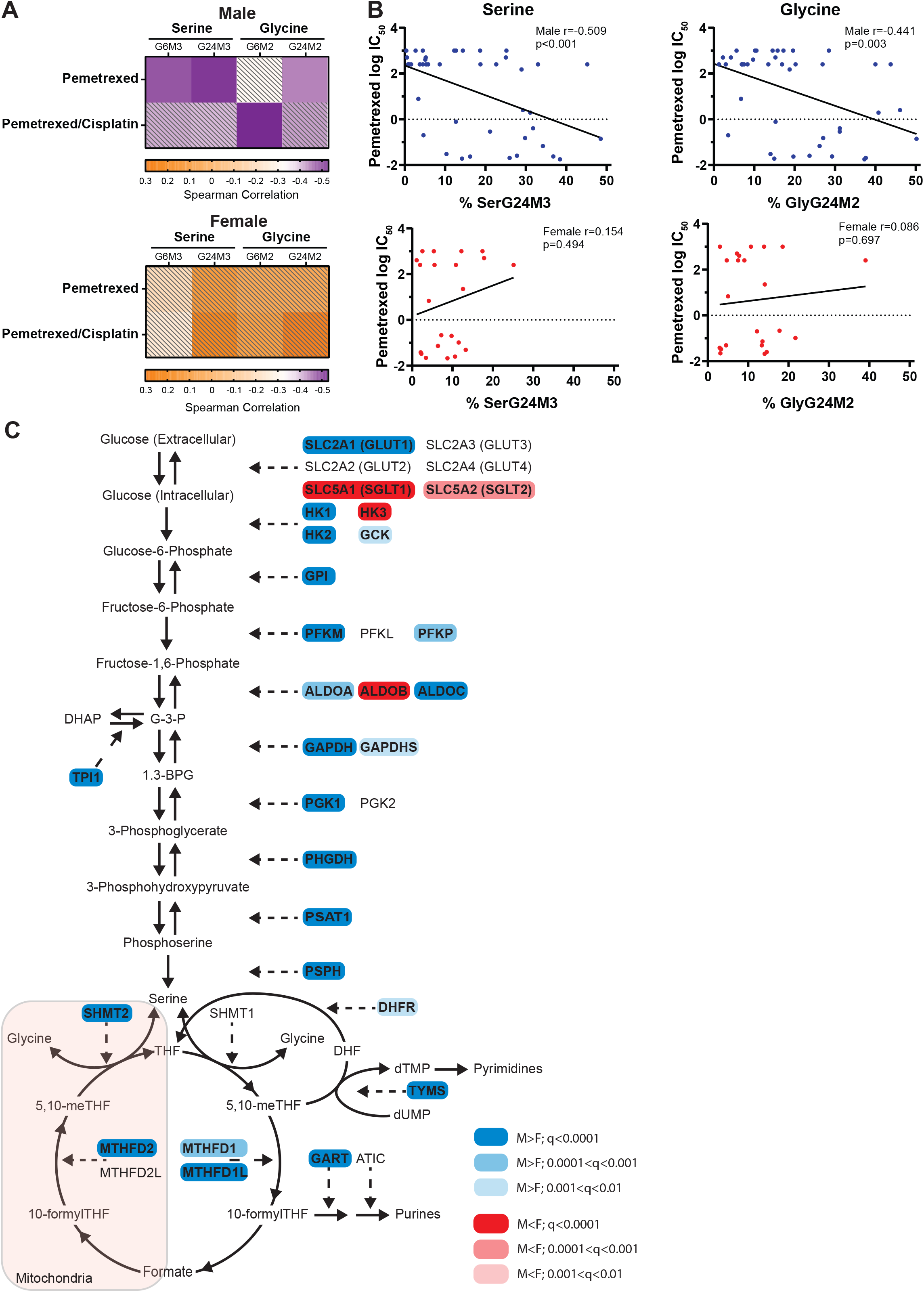
*De novo* serine biosynthesis from glucose is associated with sensitivity to antifolate drugs in male NSCLC cell lines. A, Heatmaps representing spearman correlation coefficients of [^13^C_6_] glucose incorporation into serine (m+3) and glycine (m+2) isotopologues and sensitivity to pemetrexed or pemetrexed/cisplatin in male (n=43) and female (n=22) non-small cell lung cancer (NSCLC) cell lines. Colored squares reached FDR cut-off of 0.05. B, Spearman correlation plot of [^13^C6] glucose incorporation into serine (m+3) and glycine (m+2) isotopologues and pemetrexed IC_50_ values of male (n=43) and female (n=22) NSCLC cell lines. C, Pathway schematic of enzymes involved in the *de novo* biosynthesis of serine from glucose and the folate metabolism demonstrating mRNA expression of male (n=592) and female (n=400) NSCLC tumor samples with a male (blue) or female (red) bias. Color shade represents grade of significance. **q<0.01, ***q<0.001, ****q<0.0001; Mann-Whitney test, FDR-adjusted p-values (FDR<0.01). Abbreviations: ser, serine; gly, glycine; G, glucose

Next, we examined mRNA expression levels of 35 genes involved in the *de novo* biosynthesis of serine from glucose and the folate cycle in male (n=592) and female (n=400) NSCLC samples from the TCGA Pan-Cancer dataset. The majority of the genes were expressed at higher levels in male vs female patient samples (**Fig. 2C, Fig. S1A**), including phosphoglycerate dehydrogenase (PHGDH), the rate-limiting enzyme of the serine biosynthesis pathway. This suggests that *de novo* serine biosynthesis and the folate cycle may be more important in male NSCLCs. Because pemetrexed has greater efficacy in adenocarcinomas, we assessed mRNA expression levels of the same genes in patients with adenocarcinoma and squamous cell carcinoma separately. Notably, there were fewer significantly different genes that may be due to sample size (**Fig. S2, Fig. S3**). However, of the genes that were significant, most genes were expressed at significantly greater levels in males, paralleling the NSCLC mRNA expression data. Interestingly, three folate cycle genes thymidylate synthase (TYMS; the primary target of pemetrexed (Adjei, 2004; Curtin and Hughes, 2001), methylene tetrahydrofolate reductase (MTHFD2), and 5-Aminoimidazole-4-Carboxamide Ribonucleotide Formyltransferase/IMP Cyclohydrolase (ATIC) were enriched only in male adenocarcinomas (**Fig. S2A**). Together, these data suggest that male NSCLCs may depend more on *de novo* serine biosynthesis than female NSCLCs to supply the folate cycle.

### *De novo* serine biosynthesis is associated with sensitivity to antifolate drugs in male pan-cancer

Next, we wanted to test if the sex-specific associations of pemetrexed cytotoxicity with *de novo* serine biosynthesis were generalizable to other cancers. We correlated mRNA expression levels of the genes involved in the *de novo* biosynthesis of serine from glucose and the folate cycle with drugs that inhibit the folate cycle and downstream pathways (i.e., methotrexate and 5-fluorouracil) on a cohort of pan-cancer cell lines obtained from TCLE using both area under the curve (AUC) and IC_50_ parameters derived from dose-response curves for cytotoxicity. Significant negative correlations between folate cycle gene expression and IC_50_ and AUC values (i.e., higher gene expression corresponded to lower drug dose needed for cytotoxicity), indicated that male and female pan-cancer cell lines with high folate cycle gene expression are sensitive to antifolate drugs (**Fig. 3A)**. While there was no significant difference in the number of correlations between the folate cycle genes and antifolate drug cytotoxicity in male and female cell lines, (22 significant correlations in males, 21 significant correlations in females; p=1.0, Fisher’s Exact Test), only male cell lines exhibited a significant enrichment of *de novo* serine synthetic genes whose expression correlated negatively with antifolate drug IC_50_ and AUC values (32 significant correlations in males, 17 significant correlations in females; p=0.022, Fisher’s Exact Test). This indicates that increased gene expression of *de novo* serine biosynthesis enzymes predicts sensitivity to antifolates in male pan-cancer cell lines only. The *de novo* serine synthetic genes PHGDH, PSAT1, and SHMT2 as well as other glycolytic genes such as hexokinase 2 (HK2), phosphofructokinase L (PFKL), and aldolase C (ALDOC) showed significant negative correlations between gene expression and antifolate drug IC_50_ and AUC values in male pan-cancer cell lines (**Fig. 3A**). Also of note, there were significant positive correlations between GLUT1 (SLC2A1) and hexokinase 1 (HK1) gene expression and antifolate drug cytotoxicity in both males and females, suggesting a potential mechanism for antifolate resistance. Together, these data indicate that, similar to NSCLC cell lines, *de novo* serine biosynthesis may be a predictive marker for antifolate drug sensitivity in male, but not female, pan-cancer cell lines.

**Figure 3:**
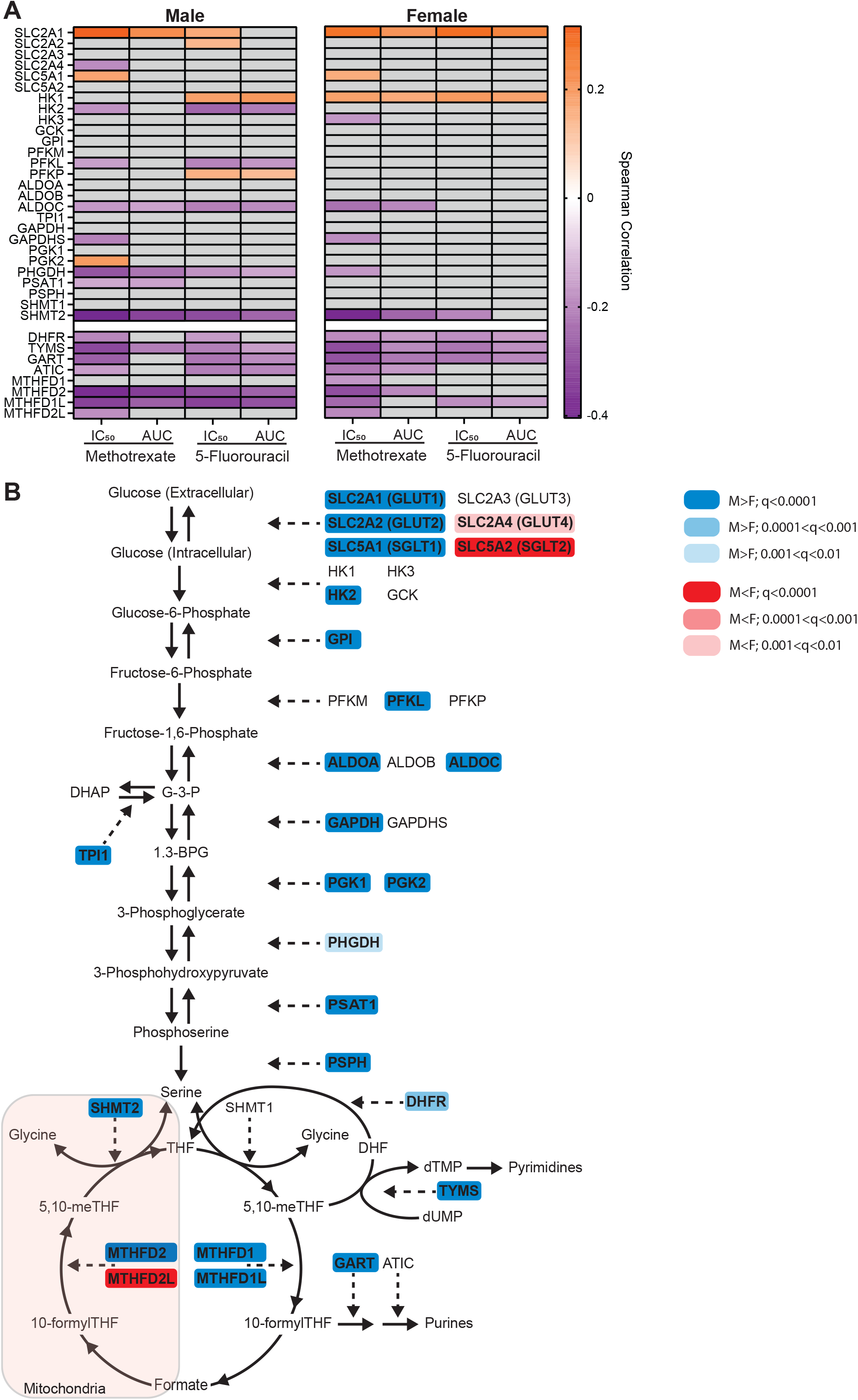
*De novo* serine biosynthesis is associated with sensitivity to antifolate drugs in male pan-cancer. A, Heatmaps representing spearman correlation coefficients of mRNA expression levels and cytotoxicity to antifolate drugs in male (n=325) and female (n=276) pan-cancer cell lines. Colored squares reached FDR cut-off of 0.05. B, Pathway schematic of enzymes involved in the *de novo* biosynthesis of serine from glucose and the folate cycle demonstrating mRNA expression of 4,076 male and 2,782 female pan-cancer tumor samples with a male (blue) or female (red) bias. Color shade represents grade of significance. **q<0.01, ***q<0.001, ****q<0.0001; Mann-Whitney test, FDR-adjusted p-values (FDR<0.01).

Next, we examined mRNA expression levels of the same genes in male (n=4,076) and female (n=2,782) tumors from the TCGA Pan-Cancer dataset. Similar to NSCLC samples, most of the genes were expressed at higher levels in male tumors (**Fig. 3B, Fig. S1B**), suggesting that the sex differences in *de novo* serine biosynthesis may extend to other cancers beyond NSCLC.

## Discussion

Targeting metabolic adaptations of cancer cells is a promising therapeutic approach. However, to this date, sex differences in cancer metabolism have not been taken much into account. Here, we report for the first time that male and female NSCLC cells exhibit different metabolic features that predict sex-biased sensitivity to cytotoxic reagents. We found that male NSCLC cell lines exhibited increased *de novo* serine and glycine biosynthesis from glucose compared to female cell lines. Furthermore, we showed that *de novo* serine biosynthesis predicts sensitivity to pemetrexed in male, but not female NSCLC cell lines, suggesting that *de novo* serine biosynthesis may be an important key player in male NSCLC. We validated these findings using transcriptome analyses of human NSCLC and pan-cancer tumors from TCGA, establishing that male tumors had increased transcript enrichment of the *de novo* serine biosynthesis pathway and the folate cycle. In addition, we showed that increased expression of serine biosynthesis genes predicted increased antifolate cytotoxicity in male, but not female, pan-cancer cell lines from TCLE.

The antifolate pemetrexed is a front-line chemotherapeutic for patients with non-squamous NSCLC (Ettinger et al., 2021). Although pemetrexed-based treatment regimens showed improved survival and improved overall response rate in males and females, about 50% of NSCLC patients continue to show no response to treatment (Ettinger, 2002; Jo et al., 2018; Park et al., 2015; Shih et al., 2020). Thus, better methods to predict therapeutic responses to pemetrexed in NSCLC patients are needed. Here, we show for the first time that stable isotope labeling studies with glucose can predict sensitivity to antifolates on a sex-specific basis *in vitro*. Thus, these data provide the basis for future research to determine whether the phosphoserine pathway may be a sex-specific biomarker for antifolate treatment response in NSCLC and other cancers.

Multiple publications support that the folate cycle may differ between males and females. Certain polymorphisms in the 5,10-methylenetetrahydrofolate reductase (MTHFR) gene, an enzyme of the folate cycle, are associated with increased or decreased risk of lung cancer in females, but not males (Shi et al., 2005), which further underlines the importance of considering sex in antifolate targeting approaches. Although the mechanisms for this clinical finding are unclear, published data indicate the presence of sex differences in the function of this pathway. In the context of normal metabolism, multiple reports have identified sex differences in folate input into one carbon metabolism. Males typically have lower folate levels, requiring increased supplementation (Cohen et al., 2021; Winkels et al., 2008). Additional data identify that abnormal folate metabolism results in sex differences in fetal development and the presence of neural tube defects (Padmanabhan et al., 2017; Poletta et al., 2018). To support these findings further, there are sex differences in the activity of one-carbon metabolism where females have higher methionine flux compared to males (Sadre-Marandi et al., 2018; Witham et al., 2013). These findings suggest a potential avenue for advancement in predicting female therapeutic response. Because *de novo* biosynthesis of serine and glycine from glucose is associated with sensitivity to antifolates only in male cell lines; female sensitivity to antifolates may be dependent upon input of other nutrients into one-carbon metabolism. However, this needs to be established.

While the data we provided here suggests that altered serine biosynthesis may sensitize male NSCLC to pemetrexed and may therefore function as a predictive marker for response to pemetrexed treatment in males, this does not necessarily suggest that male NSCLC patients respond better to pemetrexed than female patients. In fact, clinical data has shown that while pemetrexed improved survival and overall response rate in men and women (Jo et al., 2018; Park et al., 2015; Shih et al., 2020), female NSCLC patients respond better to pemetrexed-based treatment combinations (Paz-Ares et al., 2012; Wang et al., 2017). There are likely other pathways whose gene expression may correlate with antifolate response, and these pathways may be upregulated in female NSCLC. Thus, additional research is required to better understand the sex-biased outcome in male and female patient populations that received pemetrexed.

Established cancer cell lines are used in almost every aspect of basic cancer research. However, little to no attention is given to the sex of these cell lines. Here, we provide evidence that established cancer cell lines exhibit sex differences in metabolism even after years of *in vitro* culture. In addition to sex differences in *de novo* serine biosynthesis in NSCLC cell lines, we recently reported that glioma cell lines exhibit sex differences in glutaminase 1 expression (Sponagel et al., 2021), suggesting that sex differences in cancer cell lines persist across multiple metabolic pathways. Not only do these findings underline the importance of correctly identifying and considering the sex of cancer cell lines in the laboratory setting, but it identifies a completely unexplored and very important scientific space. The fact that cell lines such as these have been used for years in laboratory research and still retain sex differences in fundamental metabolic pathways suggest that the selective advantage of retaining these characteristics in cell culture are significant and outweigh the selective pressures of conventional laboratory cell culture. As the majority of genes encoding enzymes involved in central carbon metabolism are not present on sex chromosomes, these findings suggest the presence of epigenetic influences on sex differences, as has been previously identified in cancers including lung (Lopes-Ramos et al., 2018, 2020; Vaissière et al., 2009; Wu et al., 2008).

### Limitations of the study

The most important limitation of this study is that it is retrospective and limited to a subset of metabolites and nutrient utilization pathways in central carbon metabolism. Specifically, the stable isotope labeled nutrient analyzed in this study was [^13^C_6_] glucose and its analyses was restricted to serine, glycine, and selected glycolytic and TCA cycle metabolites. Emerging data from our group also showed sex differences in anaplerosis from glutamine in GBM and that these sex differences may not come from carbons as part of the TCA cycle, but rather nitrogens for transamination and amino acid synthesis (Sponagel et al., 2021). Therefore, follow up studies would need to be performed to confirm whether similar sex differences exist in NSCLC. In addition, although the use of TCLE cell lines further support the presence of sex differences in cell line and tumor metabolism and its impact on chemotherapeutic sensitivity, this is also retrospective and more work needs to be done to characterize these sex-specific metabolic-therapeutic connections on a cancer-by-cancer basis.

Furthermore, while our data provide strong *in vitro* evidence that altered *de novo* serine biosynthesis predicts sensitivity in male NSCLC and pan-cancer cell lines, additional clinical data is required to provide evidence that altered *de novo* serine biosynthesis can stratify male, but not female, patients into pemetrexed-responsive and pemetrexed-non-responsive patient cohorts.

Nonetheless, the combination of human cell line and tumor tissue data provide strong evidence to further support the emerging paradigm of factoring sex and metabolism into cancer therapeutic response. As mentioned above, the presence of sex differences in normal metabolism and the persistent sex disparity in incidence and mortality suggest that these findings may be applied to additional other cancers.

## Acknowledgements

This work was supported by the Cancer Biology Pathway Molecular Oncology Training Grant NIH T32CA113275 (JS), the NIH grants K99/R00 CA218869 (JEI), R21 CA242221 (JEI), R01 CA174737-06 (JBR), the Alvin J. Siteman Cancer Center Investment Program / The Foundation for Barnes-Jewish Hospital (JBR, JEI) and the Barnard Research Fund (JBR, JEI), and Joshua’s Great Things (JBR). This research was inspired by the Transdisciplinary Research in Energetics and Cancer (TREC) Training Workshop: R25CA203650 (PI: Melinda Irwin). The results shown here are in part based upon data generated by the TCGA Research Network: https://www.cancer.gov/tcga.

## Author Contributions

Conceptualization: JS, SD, and JEI. Investigation and formal analysis: JS, JL, and JEI performed and analyzed experiments; JS, JL, and JEI helped with data collection and analysis; Data curation: JL. Supervision: SD and JEI. Visualization: JS JL and JEI. Writing – original draft: JS and JEI. Writing – review & editing: All authors. Funding acquisition: JBR, JEI.

## Methods

### Resource Availability

#### Lead Contact

Further information and request for resources and reagents should be directed to and will be fulfilled by the lead contact Joseph E. Ippolito.

#### Materials Availability

There are no restrictions to the availability of all materials mentioned in the manuscript. Data and

#### Code Availability

Original data for Figure 1 and 2 in the paper are available in (Chen et al., 2019).

### Experimental Model and Subject Details

No experimental models or subjects were used here.

### Method Details

#### Cell line stable isotope labeling datasets

[^13^C_6_] glucose labeling data for 87 NSCLC (n=57 male and n=30 female) cell lines were downloaded as supplemental information from the publication (Chen et al., 2019). The sex of each of the cell lines was verified with the Cellosaurus database (https://web.expasy.org/cellosaurus/) (Bairoch, 2018). Isotopologue data were converted into total labeled metabolite data by summing up the percentage abundance of each of the labeled isotopologues (i.e., m + 1…m + n) for each metabolite. Total labeled metabolites were used in subsequent sex-based comparisons.

#### Sex-based correlations of drug cytotoxicity and labeled metabolites in NSCLC cell lines

Drug cytotoxicity data for the 87 NSCLC cell lines and 20 drugs were downloaded as supplemental information from the publication (Chen et al., 2019). Nonparametric Spearman correlation coefficient was calculated between each drug IC_50_ value and the total label of serine and glycine, and linear regression was fit to model the linear relationship between the IC_50_ value of pemetrexed and total label of serine and glycine. Correlations with an FDR adjusted p-value<0.05 were considered statistically significant.

#### TCGA Gene Expression Analyses

All gene expression data were obtained through cBioPortal (http://cbioportal.org). Batch normalized RNASeq RSEM data of male and female NSCLC patient samples (n=510 adenocarcinoma and n=482 squamous cell carcinomas) corresponding to 35 genes involved in *de novo* serine biosynthesis and the folate cycle were downloaded from the TCGA Pan-Cancer dataset. Adenocarcinomas included 236 male and 274 female samples and squamous cell carcinomas included 356 males and 126 females. Batch normalized RNASeq RSEM data of male and female pan-cancer patient samples corresponding to 35 genes involved in *de novo* serine biosynthesis and the folate cycle were downloaded from the TCGA Pan-Cancer dataset. It included 4,076 male and 2,782 female samples. Sex-specific cancers (i.e., breast, ovarian, endometrial, cervical, prostate, and testicular cancers) were excluded.

#### The Cancer Cell Line Encyclopedia (TCLE) Gene Expression Analyses

All TCLE data (Ghandi et al., 2019; Nusinow et al., 2020) were obtained through cBioPortal (http://cbioportal.org). The sex of each cell line was obtained from TCLE data and confirmed using the Cellosaurus database (Bairoch, 2018). Only cell lines that had the same sex assigned in both databases were included. Any cell line that was flagged by Cellosaurus as “problematic” because of contamination was excluded. The final dataset included 325 male and 267 female cell lines. Cytotoxicity data for the chemotherapeutic drugs methotrexate and 5-fluorouracil, which target the folate cycle and downstream pathways, were obtained as well as the RPKM expression analyses for 35 genes involved in *de novo* serine biosynthesis and the folate cycle. Note that pyrimethamine data were excluded, as the cytotoxicity data were absent for 65% of the cell lines. The drug sensitivity measurements were IC_50_ and “area under the curve” (AUC) values which are both derived from a sigmoidal curve by fitting a mixed-effects model (Vis et al., 2016). AUC values may be used to compare the effect of a drug across multiple cell populations in a more robust fashion than IC_50_ (Fallahi-Sichani et al., 2013). Nonparametric Spearman correlations of each drug cytotoxicity parameter were made against the expression of each of the 35 genes. Following correction of p values for FDR<0.05, only those genes that remained significant were plotted on the heatmaps. A Fisher exact test was performed to assess if there were significantly different numbers of correlations between male and female cell lines in either *de novo* serine biosynthesis or the folate cycle.

### Quantification and Statistical Analysis

#### Data representation and statistical analysis

All data except for Fig. 1B were graphed and analyzed using GraphPad Prism software version 9.0. Data for Fig. 1B were graphed using R (http::/cran.r-project.org; version 4.1.1) *ggplot2* package. All violin plots show the median and quartiles. For statistical analysis of two groups, test for normality distribution was first performed, using the D’Agostino-Pearson omnibus K2 test. If samples had a normal distribution an unpaired, two-tailed, parametric t-test was performed. If samples did not have a normal distribution an unpaired, two-tailed, non-parametric Mann-Whitney test was performed. For correlation analyses, nonparametric Spearman correlation coefficient was calculated, and linear regression was fit to model the linear relationship between two markers. Unless noted otherwise, an FDR adjusted p-value (termed q value) or raw p-value<0.05 as noted was considered statistically significant.

## Supplemental Information Titles and Legends

**Figure S1: mRNA expression of genes involved in the *de novo* biosynthesis of serine from glucose and the folate cycle in male and female NSCLC and pan-cancer tumor samples, related to Figure 2 and Figure 3**

A-B, Batch normalized RNASeq mRNA expression levels (log_2_(RSEM+1) of male (n=592) and female (n=400) NSCLC tumor samples (A) and of 4,076 male and 2,782 female pan-cancer tumor samples (B). **q<0.01, ***q<0.001, ****q<0.0001; Mann-Whitney test, FDR-adjusted p-values (FDR<0.01).

**Figure S2: Sex differences in mRNA expression levels of glycolysis, serine synthesis, and folate metabolism in adenocarcinoma and squamous cell carcinoma, related to Figure 2**

A-B, Pathway schematic of enzymes involved in the *de novo* biosynthesis of serine from glucose and the folate metabolism demonstrating mRNA expression of male (n=236) and female (n=274) adenocarcinoma patient samples (A) and male (n=356) and female (n=126) squamous carcinoma patient samples (B) with a male (blue) or female (red) bias. Color shade represents grade of significance. *q<0.05, **q<0.01, ***q<0.001, ****q<0.0001; Mann-Whitney test, FDR-adjusted p-values (FDR<0.05).

**Figure S3: mRNA expression of genes involved in the *de novo* biosynthesis of serine from glucose and the folate cycle in male and female adenocarcinoma and squamous cell carcinoma tumor samples, related to Figure 2**

A-B, Batch normalized RNASeq mRNA expression levels (log_2_(RSEM+1) of male (n=236) and female (n=274) adenocarcinoma tumor samples (A) and male (n=356) and female (n=126) squamous carcinoma tumor samples (B). *q<0.05, **q<0.01, ***q<0.001, ****q<0.0001; Mann-Whitney test, FDR-adjusted p-values (FDR<0.05).

