## Supplementary Materials for "De novo serine biosynthesis from glucose predicts sex-specific response to antifolates in non-small cell lung cancer cell lines"

Figure S1

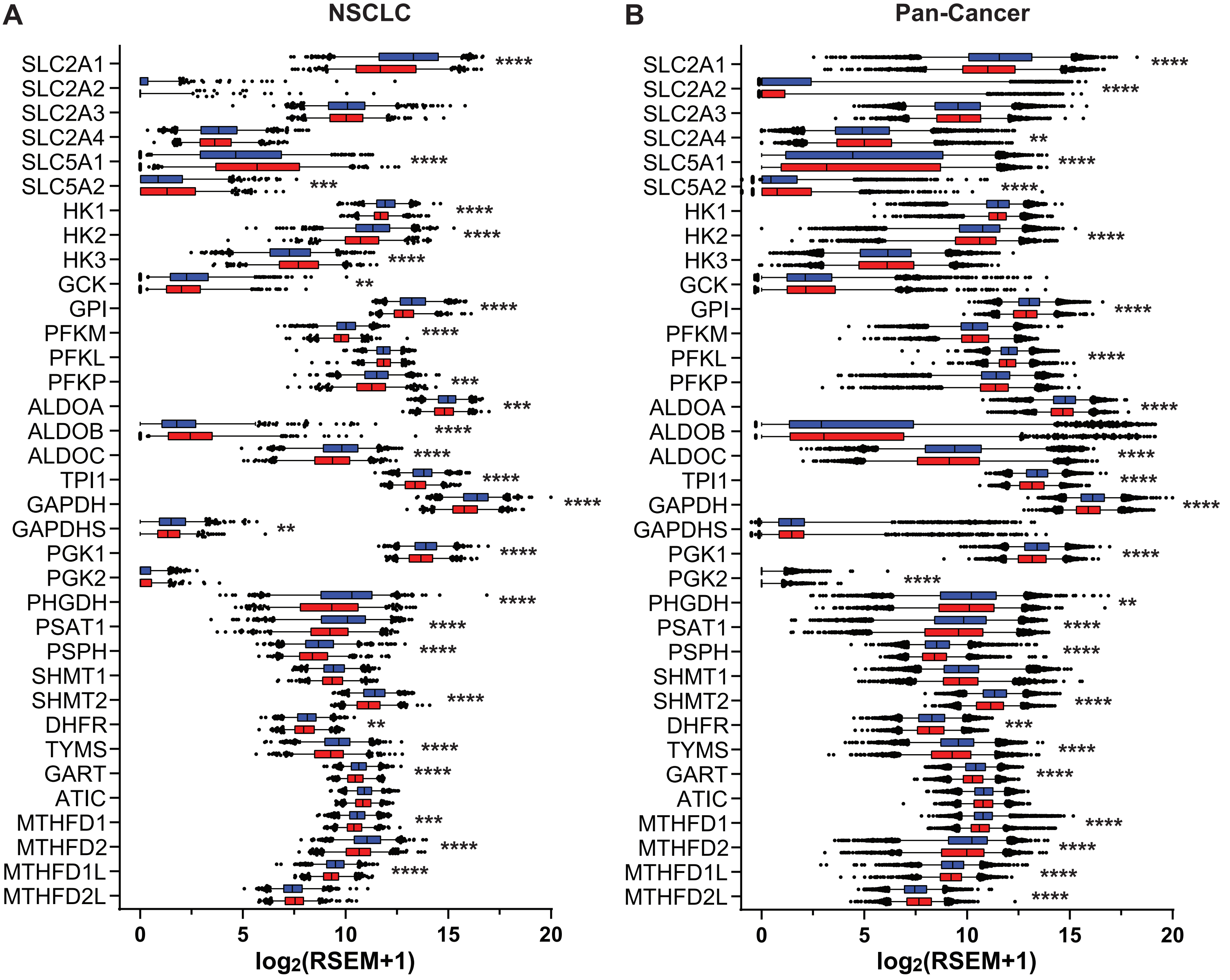
A
NSCLC
B
Pan-Cancer
SLC2A1
SLC2A2
SLC2A3
SLC2A4
SLC5A1
SLC5A2
HK1
HK2
HK3
GCK
GPI
PFKM
PFKL
PFKP
ALDOA
ALDOB
ALDOC
TPI1
GAPDH
GAPDHS
PGK1
PGK2
PHGDH
PSAT1
PSPH
SHMT1
SHMT2
DHFR
TYMS
GART
ATIC
MTHFD1
MTHFD2
MTHFD1L
MTHFD2L
0
5
10
15
20
log2(RSEM+1)

**Figure S1: mRNA expression of genes involved in the *de novo* biosynthesis of serine from glucose and the folate cycle in male and female NSCLC and pan-cancer tumor samples, related to Figure 2 and Figure 3**

A-B, Batch normalized RNASeq mRNA expression levels (log2(RSEM+1) of male (n=592) and female (n=400) NSCLC tumor samples (A) and of 4,076 male and 2,782 female pan-cancer tumor samples (B). **q<0.01, ***q<0.001, ****q<0.0001; Mann-Whitney test, FDR-adjusted p-values (FDR<0.01).

### Figure S2

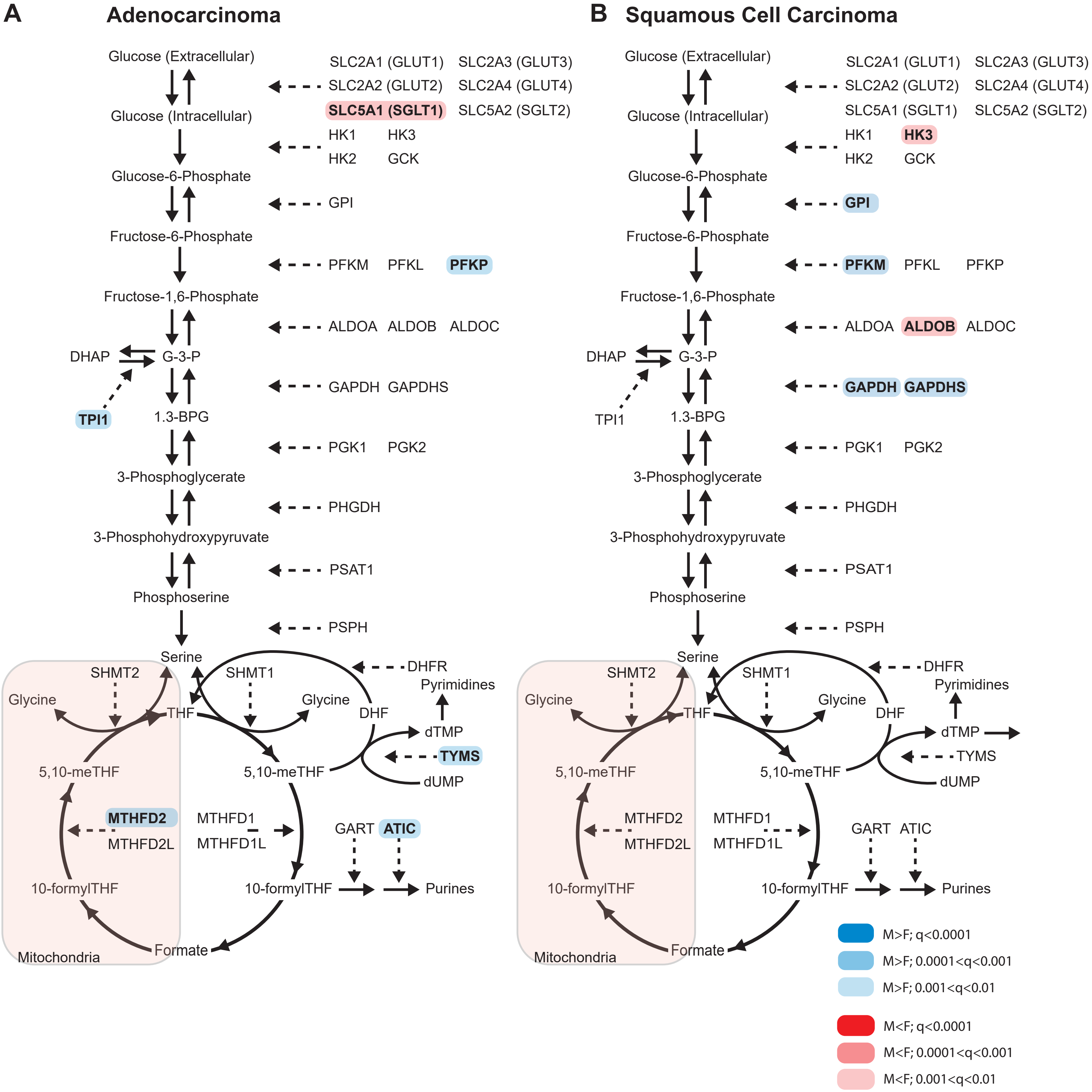
A
Adenocarcinoma
Glucose (Extracellular)
SLC2A1 (GLUT1)
SLC2A3 (GLUT3)
SLC2A2 (GLUT2)
SLC2A4 (GLUT4)
SLC5A1 (SGLT1)
SLC5A2 (SGLT2)
Glucose (Intracellular)
HK1
HK3
HK2
GCK
Glucose-6-Phosphate
GPI
Fructose-6-Phosphate
PFKM
PFKL
PFKP
Fructose-1,6-Phosphate
ALDOA
ALDOB
ALDOC
DHAP
G-3-P
GAPDH
GAPDHS
TPI1
1.3-BPG
PGK1
PGK2
3-Phosphoglycerate
PHGDH
3-Phosphohydroxypyruvate
PSAT1
Phosphoserine
PSPH
Serine
DHFR
SHMT2
SHMT1
Pyrimidines
Glycine
THF
Glycine
DHF
dTMP
TYMS
5,10-meTHF
5,10-meTHF
dUMP
MTHFD2
MTHFD1
MTHFD2L
MTHFD1L
GART
ATIC
10-formylTHF
10-formylTHF
Purines
Mitochondria
Formate
B
Squamous Cell Carcinoma
Glucose (Extracellular)
SLC2A1 (GLUT1)
SLC2A3 (GLUT3)
SLC2A2 (GLUT2)
SLC2A4 (GLUT4)
SLC5A1 (SGLT1)
SLC5A2 (SGLT2)
Glucose (Intracellular)
HK1
HK3
HK2
GCK
Glucose-6-Phosphate
GPI
Fructose-6-Phosphate
PFKM
PFKL
PFKP
Fructose-1,6-Phosphate
ALDOA
ALDOB
ALDOC
DHAP
G-3-P
GAPDH
GAPDHS
TPI1
1.3-BPG
PGK1
PGK2
3-Phosphoglycerate
PHGDH
3-Phosphohydroxypyruvate
PSAT1
Phosphoserine
PSPH
Serine
DHFR
SHMT2
SHMT1
Pyrimidines
Glycine
THF
Glycine
DHF
dTMP
TYMS
5,10-meTHF
5,10-meTHF
dUMP
MTHFD2
MTHFD1
MTHFD2L
MTHFD1L
GART
ATIC
10-formylTHF
10-formylTHF
Purines
Mitochondria
Formate
M>F; q<0.0001
M>F; 0.0001<q<0.001
M>F; 0.001<q<0.01
M<F; q<0.0001
M<F; 0.0001<q<0.001
M<F; 0.001<q<0.01

**Figure S2: Sex differences in mRNA expression levels of glycolysis, serine synthesis, and folate metabolism in adenocarcinoma and squamous cell carcinoma, related to Figure 2**

A-B, Pathway schematic of enzymes involved in the *de novo* biosynthesis of serine from glucose and the folate metabolism demonstrating mRNA expression of male (n=236) and female (n=274) adenocarcinoma patient samples (A) and male (n=356) and female (n=126) squamous carcinoma patient samples (B) with a male (blue) or female (red) bias. Color shade represents grade of significance. *q<0.05, **q<0.01, ***q<0.001, ****q<0.0001; Mann-Whitney test, FDR-adjusted p-values (FDR<0.05).

Figure S3

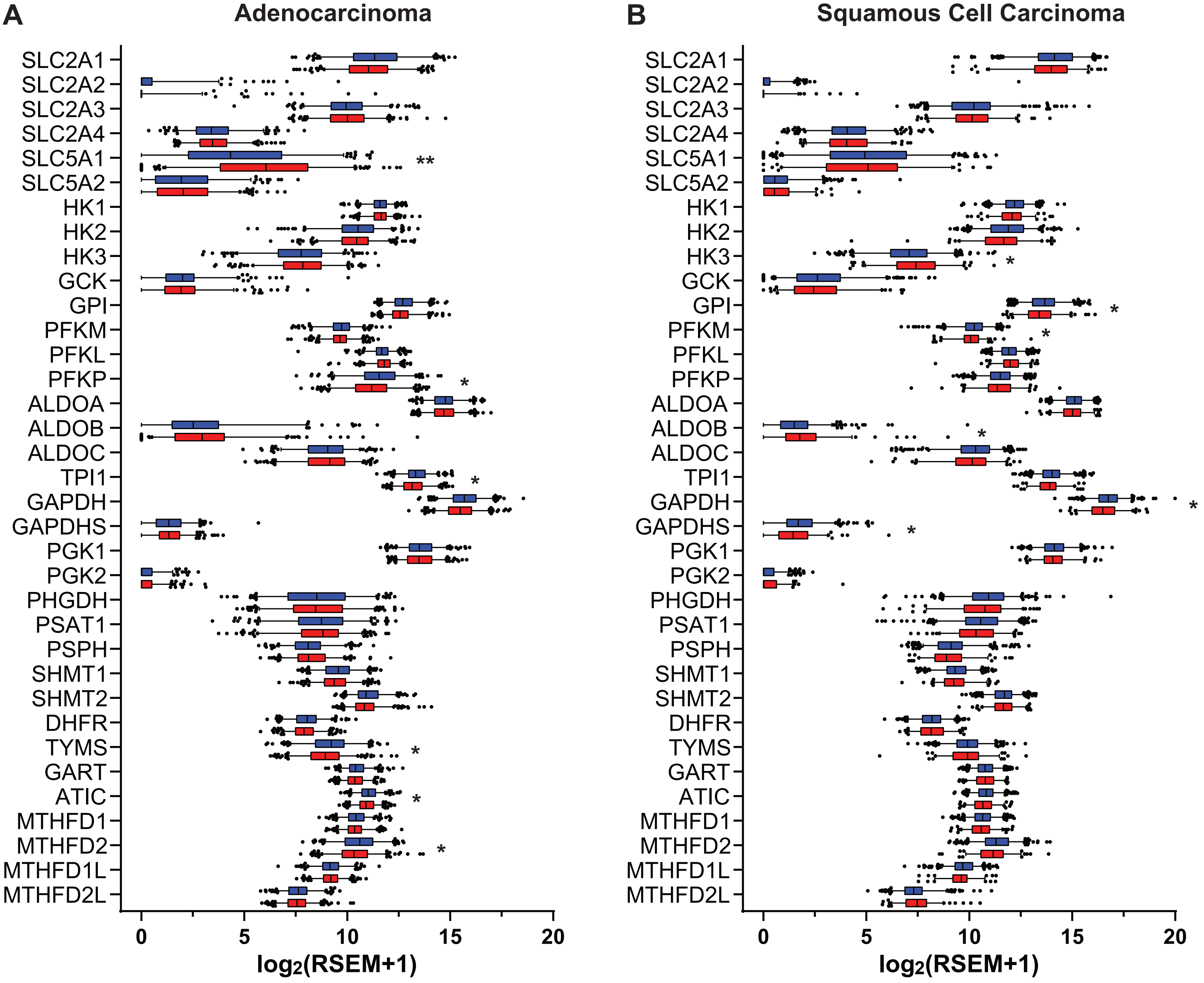
A
Adenocarcinoma
B
Squamous Cell Carcinoma
SLC2A1
SLC2A2
SLC2A3
SLC2A4
SLC5A1
SLC5A2
HK1
HK2
HK3
GCK
GPI
PFKM
PFKL
PFKP
ALDOA
ALDOB
ALDOC
TPI1
GAPDH
GAPDHS
PGK1
PGK2
PHGDH
PSAT1
PSPH
SHMT1
SHMT2
DHFR
TYMS
GART
ATIC
MTHFD1
MTHFD2
MTHFD1L
MTHFD2L
**
*
0
5
10
15
20
log2(RSEM+1)

**Figure S3: mRNA expression of genes involved in the *de novo* biosynthesis of serine from glucose and the folate cycle in male and female adenocarcinoma and squamous cell carcinoma tumor samples, related to Figure 2**

A-B, Batch normalized RNASeq mRNA expression levels (log2(RSEM+1) of male (n=236) and female (n=274) adenocarcinoma tumor samples (A) and male (n=356) and female (n=126) squamous carcinoma tumor samples (B). *q<0.05, **q<0.01, ***q<0.001, ****q<0.0001; Mann-Whitney test, FDR-adjusted p-values (FDR<0.05).
